# A simian-adenovirus-vectored rabies vaccine suitable for thermostabilisation and clinical development for low-cost single-dose pre-exposure prophylaxis

**DOI:** 10.1101/408013

**Authors:** Chuan Wang, Pawan Dulal, Xiangyang Zhou, Zhiquan Xiang, Hooman Goharriz, Ashley Banyard, Nicky Green, Livia Brunner, Roland Ventura, Nicolas Collin, Simon J Draper, Adrian VS Hill, Rebecca Ashfield, Anthony R Fooks, Hildegund C Ertl, Alexander D Douglas

## Abstract

**Background:** Estimates of current global rabies mortality range from 26,000 to 59,000 deaths per annum. Although pre-exposure prophylaxis using inactivated rabies virus vaccines (IRVs) is effective, it requires two to three doses and is regarded as being too expensive and impractical for inclusion in routine childhood immunization programmes.

**Methodology/ Principal Findings:** Here we report the development of a simian-adenovirus-vectored rabies vaccine intended to enable cost-effective population-wide pre-exposure prophylaxis against rabies. ChAdOx2 RabG uses the chimpanzee adenovirus serotype 68 (AdC68) backbone previously shown to achieve pre-exposure protection against rabies in non-human primates. ChAdOx2 differs from AdC68 in that it contains the human adenovirus serotype 5 (AdHu5) E4 orf6/7 region in place of the AdC68 equivalents, enhancing ease of manufacturing in cell lines which provide AdHu5 E1 proteins *in trans*.

We show that immunogenicity of ChAdOx2 RabG in mice is comparable to that of AdC68 RabG and other adenovirus serotypes expressing rabies virus glycoprotein. High titers of rabies virus neutralizing antibody (VNA) are elicited after a single dose. The relationship between levels of VNA activity and rabies glycoprotein monomer-binding antibody differs after immunization with adenovirus-vectored vaccines and IRV vaccines, suggesting routes to further enhancement of the efficacy of the adenovirus-vectored candidates. We also demonstrate that ChAdOx2 RabG can be thermostabilised using a low-cost method suitable for clinical bio-manufacture and ambient-temperature distribution in tropical climates. Finally, we show that a dose-sparing effect can be achieved by formulating ChAdOx2 RabG with a simple chemical adjuvant. This approach could lower the cost of ChAdOx2 RabG and other adenovirus-vectored vaccines.

**Conclusions/ Significance:** ChAdOx2 RabG may prove to be a useful tool to reduce the human rabies death toll. We have secured funding for Good Manufacturing Practice-compliant bio-manufacture and Phase I clinical trial of this candidate.

**Author summary:** Rabies was, after smallpox, the second human disease for which an efficacious vaccine was developed, by Pasteur in 1885. Although it is eminently preventable, with highly efficacious vaccines available for both humans and animals, it still causes considerable mortality in low and middle-income countries. It is a particular problem in areas with the weakest healthcare and veterinary infrastructure, where achieving prompt post-exposure vaccination or high-coverage dog vaccination are challenging.

Here, we report the development of a new candidate rabies vaccine, designed to enable low-cost single-dose pre-exposure human rabies prophylaxis in such settings. ChAdOx2 RabG is based upon a simian adenovirus-vectored candidate previously shown to achieve protection after a single dose in non-human primates, now modified to allow clinical-grade bio-manufacture. We show that it induces a potent immune response in mice, that this response can be further enhanced by clinically-relevant adjuvant, and that we can stabilise it such that it can withstand temperatures of up to 45 °C for a month. We will be performing a clinical trial of this candidate in the near future.

## Introduction

Despite the development of an efficacious rabies vaccine by Pasteur in 1885, estimates of current global annual rabies mortality range from 24,000 to 59,000 (1–3). Among the neglected tropical diseases, the burden of mortality due to rabies is exceeded only by that due to leishmaniasis (1). Across large and populous areas of Africa and Asia, rabies-attributable mortality rates exceed 1 per 100,000 people per year (3). More than 200 million individuals live in countries with rabies-attributable mortality rates exceeding 5 per 100,000 per year, corresponding to a lifetime risk of death due to rabies exceeding 0.1% (3). Such death rates exceed those attributable to some diseases included in the Expanded Programme on Immunization (EPI) and/or supported by Gavi. Calculations suggest that, in such settings, a highly-effective, simple to deliver pre-exposure prophylactic intervention costing less than USD 4 per recipient would have a cost per death averted of less than USD 4000 and a cost per DALY of less than USD 200, competitive with Gavi-funded interventions (4).

Most human cases of rabies are the result of dog bites (5). There is a strong argument for investment in dog vaccination: feasibility and cost-effectiveness of rabies control and human rabies elimination by dog vaccination has been demonstrated in some low and middle income country (LMIC) settings (6, 7). However, the countries with the highest rabies incidence are those which are least developed and most politically unstable, including Somalia, the Democratic Republic of the Congo, and Afghanistan. It remains doubtful whether adequate coverage (>60% in a canine population which turns over approximately every two years (7)) is achievable in such settings. Similarly, implementation of a robust programme of human post-exposure prophylaxis (PEP) is likely to be challenging in such settings. Following a dog bite, PEP is needed urgently and requires repeated vaccination and the administration of expensive rabies immune globulin (RIG). Achieving continuous local availability of PEP to meet such urgent yet unpredictable and intermittent demand is substantially more challenging than implementing intermittent planned mass vaccination campaigns. Concerns about the complexity of implementation of such an ‘as-needed’ intervention were the basis of Gavi’s 2013 decision not to fund PEP, despite analysis suggesting that the cost per death averted could compare favourably to that of other Gavi-funded interventions; review of this decision is expected in the near future (4).

Licensed human rabies vaccines are all based upon inactivated rabies virus (IRV), and may be used either for pre-exposure prophylaxis (PrEP) as well as PEP. PrEP regimes have until recently involved three doses of IRV vaccine spread over 28 days with typical costs of around USD 25 per vaccinee (8). The WHO recently endorsed the use of a PrEP regime involving intradermal administration of two smaller doses on each of two visits, separated by seven days: such intradermal regimes can reduce cost to USD 2 – 4 per vaccinee, but still require a total of four injections over two visits and are not licensed in some countries (9, 10). Although there are data suggesting that IRVs can be stable for a few weeks at 37 °C, their regulator-approved labels mandate refrigerated storage at 2-8 °C (11–14). As a result of these cost and delivery characteristics, rabies PrEP is not included in routine childhood immunization programmes in most rabies-endemic areas (15).

It is recommended that previous PrEP recipients who then receive a dog bite *should* still receive PEP, but this ‘post-PrEP PEP’ is an abbreviated and much cheaper course of two vaccinations without RIG. Importantly, data suggest that PrEP *alone -* without any PEP – can achieve protective antibody titers lasting many years (16, 17). This suggests that PrEP may provide substantial benefit even in contexts in which the reliable availability of PEP cannot be assured. Given the substantial proportion of children in rabies-endemic areas who receive dog bites and for whom PEP is then indicated (estimated to exceed 30% across a typical childhood in some areas (8)), it is estimated that routine PrEP could be not only more effective but *cost-saving* relative to PEP-based strategies in many contexts, particularly if the cost of PrEP is beneath USD 4 per child (8, 18).

Child-population-wide PrEP is thus an attractive intervention in areas in which Expanded Programme on Immunization (EPI) vaccines are delivered but which have otherwise limited capacity for reliable urgent PEP or for control of rabies-transmitting animals. This role for mass PrEP has been recognised, for example, in the Peruvian Amazon: in this setting, vampire bat rabies is problematic and difficult to control, access to PEP is limited, and a PrEP programme appears to have been successful (15).

Here, we have set out to develop a tool intended to enable cost-effective PrEP against rabies within routine population-wide immunization programmes. The immunological mechanism of vaccine-induced protection against rabies is well characterised. A virus neutralizing antibody (VNA) titer exceeding 0.5 international units per milliliter (IU/mL) is accepted as a marker of adequate immunization (19). This threshold is widely thought to signify clinical protection: indeed, in animal challenge studies, 100% protection is achieved at 0.1-0.2 IU/mL, with incomplete but substantial protection at even lower titers (20). Simian adenovirus-vectored vaccines are an attractive platform technology for induction of antibody responses, circumventing the problem of pre-existing anti-vector antibody to human adenovirus serotypes and readily manufacturable at large scale and low cost (21, 22). A chimpanzee adenovirus serotype 68 (AdC68)-vectored rabies vaccine and the ability of a single low dose of this vaccine to achieve long-lasting protection against rabies challenge in non-human macaques has previously been reported (23, 24). We now describe the development of a closely-related simian-adenovirus-vectored rabies vaccine, ChAdOx2 RabG, which is suitable for good manufacturing practice (GMP)-compliant production. We also describe additional approaches which may make this particularly suitable for use in low-income settings, namely thermostabilisation of the vaccine and dose-sparing adjuvantation. ChAdOx2 RabG may prove to be suitable for PrEP in highly rabies-endemic settings which have adequate infrastructure to achieve appreciable levels of childhood immunization coverage (25) but which currently lack capacity for reliable dog vaccination or PEP.

## Methods

### Plasmid and adenovirus production

Plasmid pC68 010-Rabgp comprising the E1- and E3-deleted AdC68 genome with the full-length ERA strain rabies glycoprotein coding sequence under a human cytomegalovirus immediate-early (HCMV-IE) promoter was constructed based on a virus obtained from ATCC (ATCC VR-594, Genbank accession: FJ025918.1). The wildtype AdC68 was propagated in HEK 293 cells and purified by CsCl gradient centrifugation, followed by viral genomic DNA purification as described. To generate the E1-deleted AdC68 molecular clone, the 5’ right inverted terminal repeat (ITR) was amplified by PCR and cloned into the pNEB193 vector. Using restriction enzyme sites that are unique in assembly but not necessarily unique to the full AdC68 genome, approximately 2.6 Kb of the E1 region between SnaBI and NdeI sites (from 455bp to 3028bp) were removed and replaced with a linker which contains the rare enzyme sites of I-CeuI and PI-SceI. The resultant was the pC68 000 plasmid. To delete the E3 domain, a 3.6 kb fragment was excised using AvrII and NruI (from 27793bp to 31409bp): briefly, the pC68 000 was digested by AvrII, the 5.8kb fragment was subcloned into a pUC19-like backbone (generating pXY-AvrII), and NruI was used to excise a 1.4 kb fragment (generating pXY-E3 deleted). Later, pXY-E3 deleted (insert donor) was digested with AvrII and SpeI and the insert was ligated into pAdC68 000, to produce plamid pC68 010. The HCMV-IE promoter – rabies glycoprotein cassette was inserted as previously reported (23), generating pC68 010-Rabgp (differing from previously published constructs, notably in the deletion of the E3 region).

To construct a vector for transient mammalian expression of rabies glycoprotein and as a precursor to adenovirus vector production, the full-length coding sequence was PCR amplified from pC68 010-Rabgp using oligonucleotides providing flanking Acc65I (5’) and NotI (3’) restriction enzyme sites. This permitted restriction-enzyme mediated cloning of the PCR product into pENTR4 LPTOS, a plasmid providing HCMV-IE promoter with intron A and tetracycline operator elements (26, 27). This transgene is referred to henceforth as SP_rab_-G_native,_ or simply ‘G’.

A codon-optimized version of the ERA strain glycoprotein gene in which the viral signal peptide was replaced by that of the human tissue plasminogen activator (tPA) was synthesized by ThermoFisher and cloned into pENTR4 LPTOS similarly; this transgene is referred to henceforth as SP_Tpa_-G_opt._ A third pENTR4 LPTOS plasmid in which the codon-optimized gene was preceded by the viral signal peptide was generated by InFusion cloning (Takara); this transgene is referred to henceforth as SP_rab_-G_opt_.

To produce the ChAdOx2 RabG adenoviral vector, Gateway LR recombination (ThermoFisher) was then used to transfer the SP_rab_-G_native_ transgene cassette into the ChAdOx2 parent bacterial artificial chromosome (BAC) (21). Adenoviral destination vectors for the expression of RabG by AdHu5, chimpanzee adenovirus serotype 63 (ChAd63) and ChAdOx1 were produced similarly, using previously described viral backbones (28, 29) and a full-length non-codon-optimized glycoprotein gene, with the exception that the transgene used was derived from the SAD B19 strain (Addgene plasmid 15785, a kind gift of Miguel Sena-Esteves (30)). SAD B19 and ERA were both derived from the Street Alabama Dufferin (SAD) isolate; their glycoprotein genes differ at only 4 of 524 amino acid loci (31).

AdHu5, AdC68-010, ChAd63, ChAdOx1, and ChAdOx2 adenoviruses were produced from the plasmids/ BACs described above (and subsequently titered) by the Jenner Institute Viral Vector Core Facility, as previously described (26).

Prior to genetic stability studies, ChAdOx2 RabG was subjected to two rounds of plaque picking. Virus was then propagated on adherent HEK293 cells (Oxford Clinical Biomanufacturing Facility proprietary cell bank) for five passages, prior to caesium chloride (CsCl) purification, phenol-chloroform DNA extraction and enzymatic restriction analysis, as previously described (32).

### Assessment of G expression in transiently transfected cells

G expression from the three ERA G transgene constructs in mammalian cells was assessed by flow cytometry. Two or three independent DNA preparations of each pENTR4 LPTOS plasmid were produced and transfected into HEK293E cells (National Research Council, Canada) using 25kDa linear polyethylene-imine (Polysciences), as previously described (33). Identical cells were incubated either with serum from AdHu5 RabG-immunized mice, or with naïve mouse serum (negative control), and then with Alexa488-conjugated goat-anti-mouse secondary antibody (ThermoFisher). The ratio of median fluorescence intensity (MFI) in fully-stained versus negative controls was used to quantify glycoprotein expression.

Expression of codon-optimized G constructs (G_opt_, i.e. SP_tPA_-G_opt_ and SP_rab_-G_opt_) was compared to that of the base-case construct (SP_rab_-G_native_) by dividing the G_opt_ MFI ratio by that obtained in the same experiment with SP_rab_-G_native_.

### Ethics statement

All animal work was performed in accordance with the U.K. Animals (Scientific Procedures) Act 1986, and was approved by the University of Oxford Animal Welfare and Ethical Review Body (in its review of the application for the U.K. Home Office Project Licenses PPL 30/2889 and P9804B4F1).

### Animals, vaccine preparation and immunization

Female CD-1 outbred mice (Harlan, Charles River and Envigo) were used throughout. Mice were housed in a specific-pathogen-free facility and were 6-7 weeks old at the initiation of each experiment.

All adenovirus vaccine doses were calculated on the basis of infective unit (IU) titers, but viral particle (VP) titers and hence particle: infectivity (P:I) ratios were also measured. In viral preparations used for mouse immunization, P:I ratios were 23 for AdHu5, 132 for ChAd63, 65 for ChAdOx1, 63 for AdC68, and 173 for ChAdOx2.

Addavax™ was purchased (Invivogen). SWE, a non-branded squalene-in-water emulsion was prepared at the Vaccine Formulation Laboratory in Lausanne, using a GMP-compatible manufacturing process as previously described (34). Both adjuvants were used at a dose of 25 μL per mouse, mixed by vortexing for two seconds with the appropriate adenovirus dose (diluted to 25 μL in phosphate buffered saline [PBS]) 1-2 hours prior to administration.

As comparators, we used two IRV vaccines. Rabipur™ (Novartis) consists of a liquid formulation of unadjuvanted inactivated Flury LEP strain virus, produced on purified chick embryo cells (PCEC), licensed for human use, and with a potency of >2.5 IU/mL in the NIH mouse potency assay. Nobivac Rabies™ (MSD Animal Health) consists of a liquid formulation of aluminium phosphate adjuvanted inactivated Pasteur strain virus, produced on BHK-21 cells, licensed for animal use, and with a potency of >2 IU/mL.

Vaccine doses used for each experiment are set out in the corresponding figure legends. All vaccinations were diluted in PBS to a total volume of 50 μL (with the exception of the highest doses of Rabipur™ and Nobivac Rabies™ [Figure 2] which were administered undiluted in 100 μL). Vaccine was administered intramuscularly, split equally between the gastrocnemius muscles of each hind limb.

### Virus neutralizing antibody (VNA) assay

Coded serum samples were assayed for VNA.

For data shown in Figure 2, the fluorescent antibody virus neutralization (FAVN) assay was performed at the Animal and Plant Health Agency (APHA) as previously described, with quadruplicate serum dilutions and pre-determined acceptance criteria for virus dose as measured by back titration (35).

For data shown in Figure 3, the rapid fluorescent focus inhibition test (RFFIT) was performed at the Wistar Institute, again as previously described (36), using MNA cells.

Both assays used the CVS-11 reference virus strain and the WHO international reference standard to derive titers expressed in IU/mL. The two methods are known to correlate closely (35).

### ELISA

To produce soluble rabies glycoprotein (RabG_sol_) for ELISA plate coating, the sequence encoding amino acids 1-453 of the SAD B19 strain G protein (MVPQ…DLGL) was PCR amplified from Addgene plasmid 15785 (as above). The primers used added a 5’ Acc65I site and 3’ sequence encoding a C-tag and NotI site (37), enabling cloning into pENTR4 LPTOS (as described above for adenovirus generation). This plasmid was then transfected into Expi293 cells using Expifectamine (both from ThermoFisher), in accordance with the manufacturer’s instructions. RabGsol protein was purified using a CaptureSelect C-tag column (ThermoFisher).

ELISA was performed as described previously (38). In brief, plates were coated with RabG_sol_ protein (100 ng/well in 50 μL PBS). Dilutions of test sera (in triplicate) and a standard curve produced using serial dilutions of an in-house reference serum pool from mice immunized with AdHu5 RabG, were added to the plate. Washing, secondary Ab binding, final washing, and detection were all as previously described (38), with the exception that the secondary Ab used was alkaline-phosphatase–conjugated goat anti-mouse IgG (Sigma-Aldrich). OD_405_ was quantified using ELx800 or Clariostar plate readers (Bio-Tek and BMG respectively). Results were expressed in arbitrary antibody units (AU), defined using the in-house reference standard, by interpolation of OD_405_ readings on the standard curve. Negative control sera (from mice immunised with AdHu5 expressing ovalbumin(39)) gave no detectable response.

### Thermostabilisation

Sugar-matrix thermostabilisation (SMT) was performed essentially as previously described (40). In brief, adenovirus was formulated in thermostabilisation buffer (0.4M trehalose, 0.1M sucrose). Virus was applied to Whatman Standard-14 paper (GE Healthcare) at a ratio of 50 μL/cm^2^. The virus-loaded paper was then dried for 48 hours in a glovebox (Coy Laboratory Products) at room temperature (24+/- 2 °C) and controlled humidity (% relative humidity <5%) before being transferred to airtight vials.

Moisture content of the dried product was measured by Karl-Fischer analysis using an 851 Titrando coulometer equipped with 860 Thermoprep oven (Metrohm), in accordance with the manufacturer’s instructions. The mass of water measured per cm^2^ of product was used to estimate percentage moisture content of the sugar glass, based upon the calculated 9.3mg mass of solute present in 50 μL of the thermostabilisation buffer.

As a comparator for the stability of SMT product, virus was formulated in the liquid buffer ‘A438’ (10 mM Histidine, 7.5% sucrose, 35 mM NaCl, 1 mM MgCl_2_, 0.1% PS-80, 0.1 mM EDTA, 0.5% (v/v) Ethanol pH 6.6) (41).

Thermostabilised vaccine was reconstituted by the addition of 500 μL/cm^2^ of phosphate buffered saline and vortexing for 2 seconds. The IU titer of recovered virus was measured as described for adenovirus production.

### Data analysis

Prism 7 software (Graphpad) was used for data analysis and production of graphs. Statistical analyses are described in full where reported in the Results section and Figure legends. All analyses used log_10_-transformed ELISA and VNA data; where negative ELISA results were obtained, they were assigned an arbitrary value of 3 AU (just below the detection limit) to permit log10 transformation and analysis.

## Results

### ChAdOx2 RabG is a simian adenovirus-vectored rabies vaccine suitable for GMP manufacturing and clinical development

A promising AdC68-vectored rabies vaccine has previously been described (23, 42). ChAdOx2 RabG differs from the previously reported vaccine in the following respects:

1. In addition to deletion of the E1 region, the E3 region of the adenovirus has been deleted. The E3 region is non-essential *in vitro* but encodes protein products involved in subversion of the host immune response and viral persistence (43). Its deletion is customary in the majority of adenoviruses used in clinical studies and will strengthen confidence in the relevance to the current vaccine of the growing body of literature regarding safety of E1-and E3-deleted adenovirus vectors.
2. The promoter used is the Intron A-containing HCMV-IE promoter which we have previously reported to enhance immunogenicity relative to a non-intron containing version of the promoter (27).
3. The promoter includes tetracycline operator elements, resulting in repression of transgene expression in cell lines expressing the tetracycline repressor protein (32). The use of such ‘tet-repressing’ cell lines in some GMP manufacturing facilities minimises transgene-induced effects upon adenoviral growth, enhancing the predictability of viral growth characteristics and minimising selective pressure for the outgrowth of mutant viruses (32).
4. The ChAdOx2 vector backbone contains the AdHu5 E4 orf6/7 region in place of the AdC68 equivalents. Adenoviral E4 orf6 protein forms a complex with the E1B 55K protein; the complex has multiple functions in viral growth (44). Replacement of non-AdHu5 vectors’ E4 orf6 sequence with the AdHu5 equivalent has previously been shown to enhance viral yield in cell lines supplying AdHu5 E1 proteins (29, 45). We have previously reported that ChAdOx2 virus yields are 2 to 10-fold higher than those with AdC68 (21).

We also explored the possibility of modifying the RabG transgene by codon optimisation for mammalian cells and the incorporation of the human tissue plasminogen activator signal peptide. We have previously observed enhanced levels of transgene expression and immunogenicity upon making similar changes to transgenes in other adenovirus vectored vaccines. In the case of RabG, we observed no beneficial impact of such changes upon transgene expression (Figure 1A-B). ChAdOx2 RabG thus uses the unmodified transgene (SP_rab_-G_native_, henceforth simply ‘G’).

**Figure 1:**
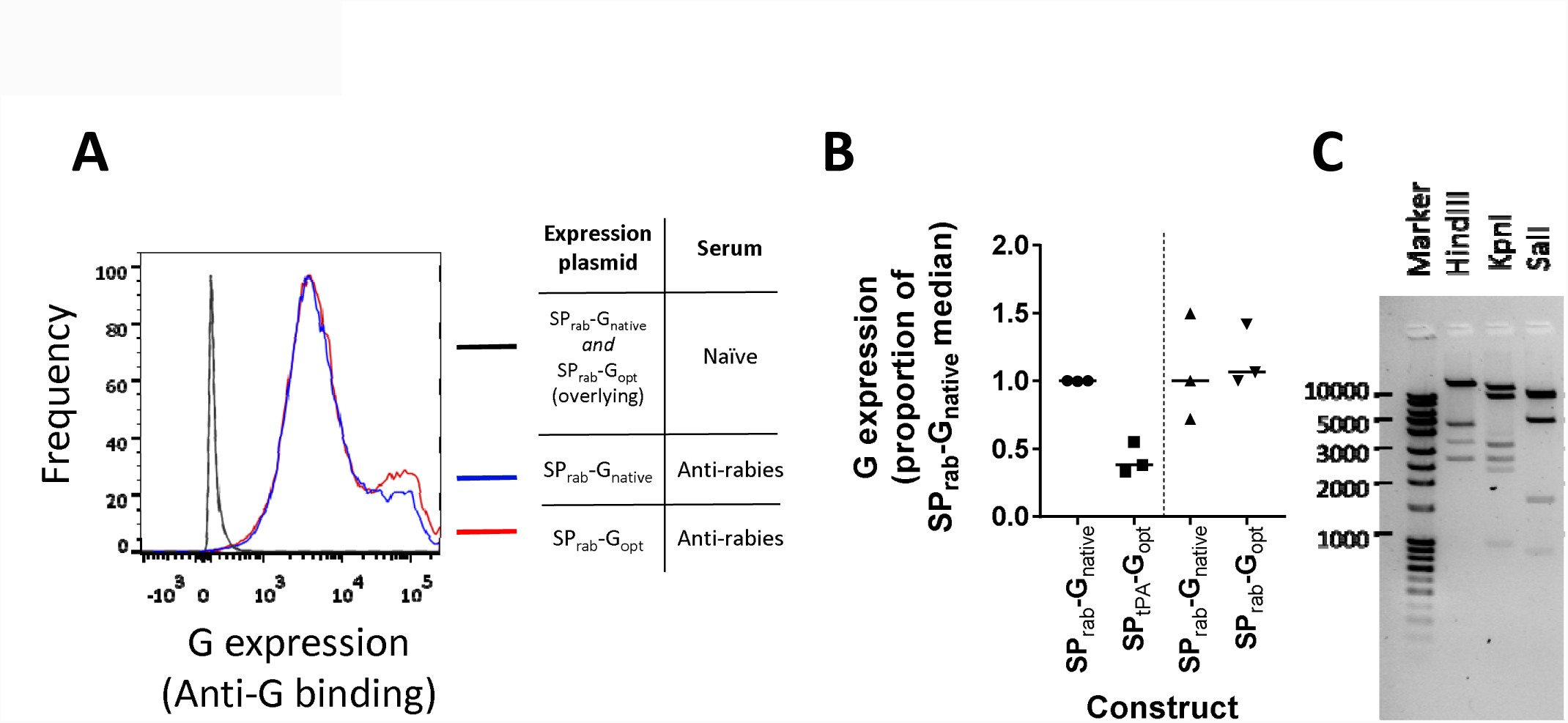
Choice of G transgene construct and demonstration of genetic stability. Panel A shows level of cell-surface expression of rabies glycoprotein after transient transfection of cells with the SP_rab_-G_native_ and SP_rab_-G_opt_ expression plasmids; legend indicates the combination of plasmid and staining conditions indicated by each line. Histograms are shown to illustrate the derivation of the data represented in Panel B: those shown here are the median results for each plasmid of three transfections with independent DNA preparations. Panel B summarizes G expression levels from the three tested constructs. Each point represents an independent transfection using an independent DNA preparation; line indicates median for each construct. SP_tPA_-G_opt_ and SP_rab_-G_opt_ were tested in separate experiments, each including the SP_rab_-G_native_ as a comparator. Because absolute fluorescence data are not comparable between experiments (fluorescence intensity varies between experiments despite using the same conditions), expression was calculated using MFIs, normalized to the SP_rab_-G_native_ MFI used in each experiment (see Methods), Separate SP_rab_-G_native_ data are thus shown for each comparison, and similarly, for clarity, SPtPA-Gopt data is not shown in panel A. Panel C shows the banding pattern of restriction-enzyme digested ChAdOx2 RabG DNA following the five-passage genetic stability study. Banding patterns were as expected. With Hind III: bands of 21974, 4451, 3349, 2645 base pairs (bp) were seen; a predicted 562bp band was faintly visible but not seen here; a predicted 96bp fragment was, as expected, not visible. With KpnI: bands of 14510, 9370, 3223, 2644, 2329 and 960bp were seen. With SalI: pairs of fragments of 10711 and 9563bp and of 4845 and 4787bp were seen as doublets; fragments of 1656 and 897bp were seen individually; a predicted 474bp band was faintly visible but not seen here; and a predicted 145bp fragment was, as expected, not visible.

To ensure suitability for manufacture in non-transgene-repressing cells (which are often used for GMP manufacture but in which production of some adenoviruses is frequently problematic (32)), we assessed virus yield in HEK293A cells (ATCC). In two independent preparations results were as follows:

1.Yield of 5.0×10^12^ VP / 9.0×10^10^ IU (P:I ratio = 54) from 5×10^7^ cells

2.Yield of 1.9×10^13^ VP / 2.1×10^11^ IU (P:I ratio = 92) from 1.5×10^8^ cells

Yield was thus approximately 1×10^5^ VP per cell, favourably comparable to our experience with other viruses (even in transgene-repressing cells), and with yields reported in the literature for other adenoviruses (46).

Following 5 passages in HEK293A cells, CsCl purification and DNA extraction, enzymatic restriction analysis demonstrated a banding pattern consistent with the starting virus sequence (Figure 1C). Although this clearly does not rule out point mutations, it does confirm genetic stability of the vector to the level required for GMP-compliant manufacture for early phase clinical trials.

### ChAdOx2 RabG elicits rabies virus neutralizing antibody in mice

To assess the immunogenicity of our adenovirus-vectored vaccine candidates, we immunized mice with one of a range of adenovirus serotypes expressing rabies glycoprotein, or one of the licenced IRVs Rabipur™ (unadjuvanted) and Nobivac Rabies™ (alum-adjuvanted). For each vaccine, dose-response was assessed. In the case of the adenoviruses, the highest dose tested was 1×10^8^ IU (c. 5×10^9^ VP, dependent upon P:I ratio). This is around 1/10 of a typical human dose of 5×10^10^ VP. In the case of the IRVs, the highest dose tested was similarly 1/10 of the manufacturer’s recommended human or dog/cat dose, i.e. >0.25 IU for Rabipur and >0.2 IU for Nobivac: this was the highest dose which could be given within the constraints of a 100 μL intramuscular injection volume.

Vaccines were assessed in two groups in separate experiments. In an initial exploratory experiment, (prior to the availability of the ChAdOx2 vector), we assessed AdHu5, ChAd63, ChAdOx1 and the IRVs. Induction of RabG_sol_-binding antibody, as measured by ELISA, was higher for the adenovirus vectors than for the IRVs, regardless of the dose used (though it should be noted that all vaccines were given as a single dose rather than the repeated dosing used for IRVs in humans) (Figure 2A). While responses to the IRVs did not rise beyond the initial timepoint (4 weeks), the adenovirus-vectored vaccines induced antibody responses which gradually rose over 12 weeks (Figure 2B). We previously observed a similar kinetic with AdC68.rabgp in macaques (23). In this initial experiment, VNA was assessed at week 12 for the highest-dose groups. All vaccines induced high VNA titers (Figure 2C). Interestingly, the relationship between RabG_sol_-binding antibody ELISA response and VNA titer was markedly different for adenovirus-vectored vaccines and IRVs (Figure 2D).

**Figure 2:**
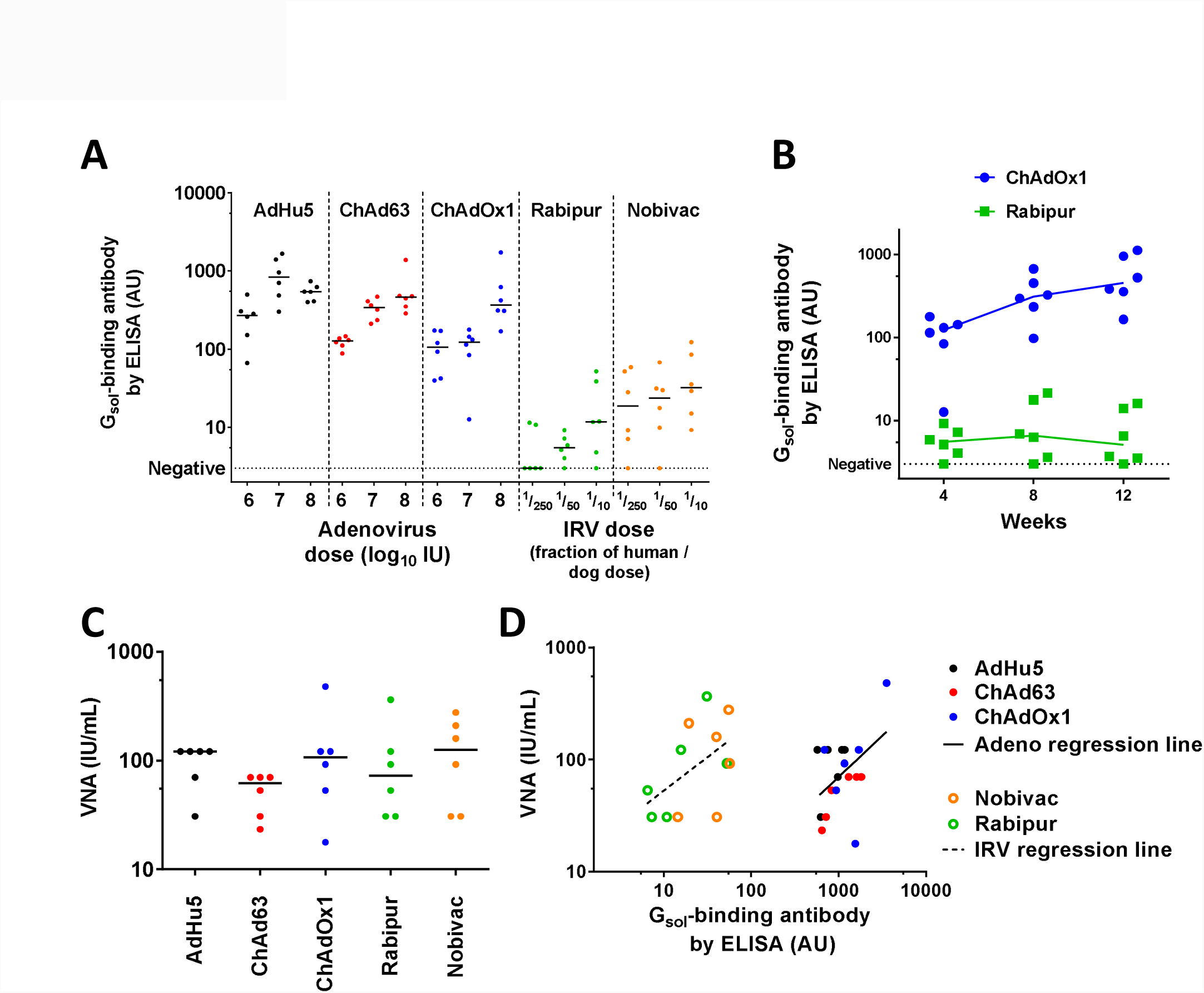
Immunogenicity of first-generation adenovirus-vectored rabies vaccines and IRVs. In all panels, points indicate results from individual mice; lines in panels A-C indicate/connect group medians. Panel A shows ELISA-measured antibody induction at 4 weeks post-immunization by AdHu5, ChAd63, and ChAdOx1 adenovirus-vectored rabies vaccines as compared to Rabipur and Nobivac IRVs. Panel B shows kinetic of antibody responses induced by ChAdOX1 (1×10^7^ IU dose group) and Rabipur (1/50 recommended human dose group); similar kinetics were seen for adenovirus-vectored vaccines and IRVs respectively, across different doses and vaccine subtypes (data not shown). For paired two-tailed t-test comparing week 4 and week 12 responses, p=0.002 for ChAdOX1; p=0.58 for Rabipur. Panel C shows VNA titers measured from week 12 samples in the groups receiving the highest doses of each vaccine (1×10^8^ IU i.e. c. 1/10 typical human dose for adenovirus-vectored vaccines, 1/10 human/dog dose for IRVs). Panel D shows relationship between ELISA-measured antibody levels and VNA titers. Filled symbols indicate adenovirus-vaccinated mice; open symbols indicate IRV-vaccinated mice. Solid line indicates fitted linear regression trend-line for adenovirus-vaccinated mice; dashed line indicates fitted regression trend-line for IRV-vaccinated mice. p=0.008 for identical intercept of the two lines, analysed by ANCOVA (no significant difference in slopes [p=0.65]).

In a subsequent experiment, we compared the immunogenicity of the ChAdOx2 RabG vector with AdC68.010.rabgp. Differences between the ChAdOx2 RabG vector, the previously reported AdC68.rabgp vector (24) and AdC68.010.rabgp are described above; the removal of E3 from AdC68.010.rabgp is not expected to affect immunogenicity. Mice receiving 1×10^7^ IU ChAdOx1 RabG were included as a group for bridging/ comparison with the previous experiment. ELISA and VNA responses induced by ChAdOx2 RabG were significantly higher than those induced by AdC68.010.rabgp (Figure 3). Interestingly the dose-response relationships differed markedly, with similar responses to the two vectors at high dose (1×10^8^ IU), but substantially stronger responses to ChAdOx2 than AdC68.010 at lower doses.

**Figure 3:**
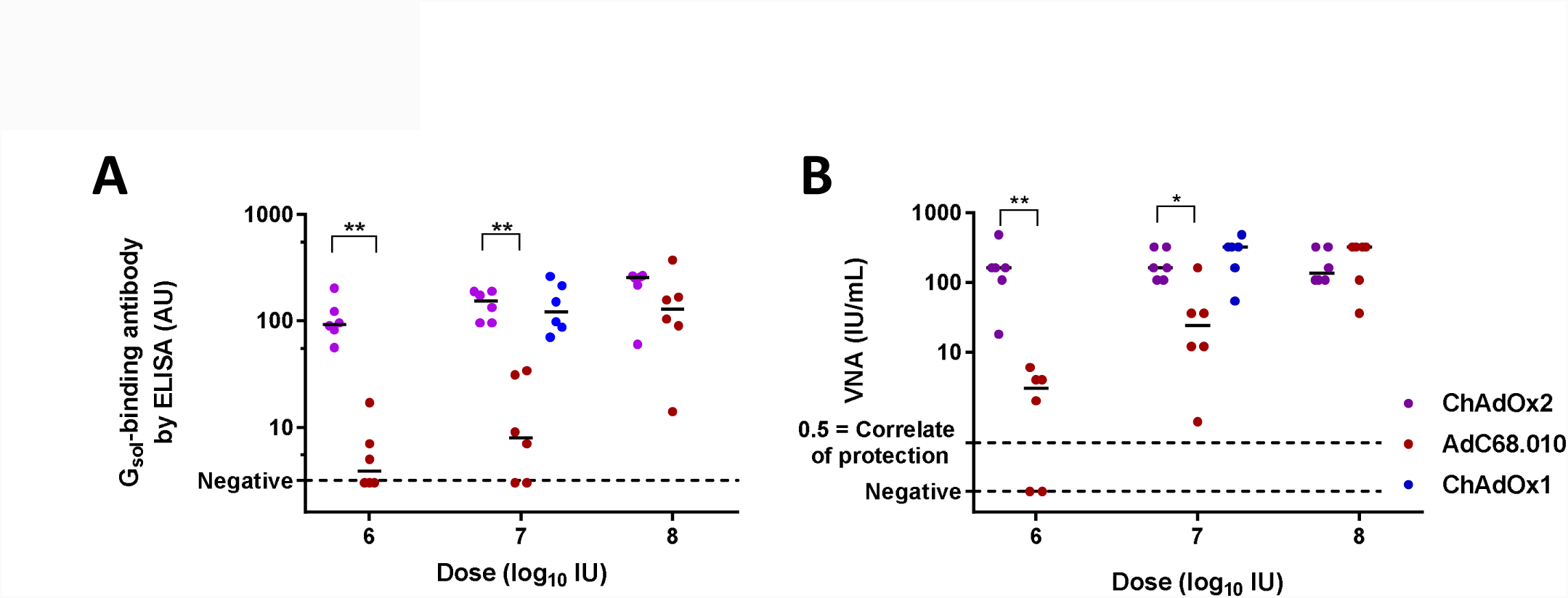
Immunogenicity of ChAdOx2 and AdC68.010 adenovirus-vectored vaccines. Panel A shows ELISA-measured antibody induction by ChAdOX2, AdC68.010 and ChAdOX1 adenovirus-vectored rabies vaccines. ChAdOX2 and AdC68.010 vaccines were given at a range of doses, as indicated on the X-axis. Serum was collected four weeks after immunization. Panel B shows VNA results for the same samples shown in panel A. In both panels, points indicate results from individual mice; lines indicate group medians.** indicates p=0.002, * indicates p=0.01 for comparisons of ELISA and VNA responses by two-tailed Mann Whitney test.

### A dose-sparing effect can be achieved by formulation of adenovirus-vectored rabies vaccines with a chemical adjuvant

Chemical adjuvants are widely used with protein and killed-virus vaccines, both to induce stronger immune responses and to achieve an antigen-dose-sparing effect. The cost-of-goods of adenovirus-vectored vaccines is likely to be relatively low (as a result of the development of high-yielding scalable manufacturing processes). Nonetheless, any reduction in the adenovirus dose required to achieve a protective response would be valuable as it could enhance the cost-efficacy of population-wide vaccination campaigns in low-income settings. We therefore sought to explore whether responses to our adenovirus-vectored rabies vaccines could be enhanced using a chemical adjuvant. We have previously reported that co-administration of adenovirus-vectored vaccines with certain adjuvants could enhance CD8^+^ T cell responses but, in contrast with others’ observations, we had not seen such effects upon antibody responses to viral vectors in the absence of co-administered protein antigen (39, 47, 48).

Here, we focussed upon squalene oil-in-water adjuvants which, to our knowledge, have not previously been explored in combination with adenovirus-vectored vaccines. In particular, we evaluated Addavax™ (Invivogen) and SWE (produced at VFL, Lausanne, Switzerland): both of these have a composition similar to MF59 (previously developed by Novartis and now marketed by GSK), which has been given to millions of people in licensed vaccines and has an excellent safety record (49). We observed a beneficial impact of such squalene emulsion adjuvants upon immunogenicity of our vaccines in each of three independent experiments, together encompassing ChAdOx1, ChAdOx2, Addavax™ and SWE (Figure 4). The enhancement in ELISA response at doses ≥1×10^6^ IU was small (statistically significant increases of c. 2-fold in antibody titer in two experiments-Figure 4A-B; not detectable in one of the three experiments-Figure 4C, right-hand side). A more marked benefit of adjuvant was apparent when very low doses of vaccine (≤5×10^4^ IU) were used (Figure 4C, left-hand side and Figure 4D); remarkably, in the presence of adjuvant, 11/12 mice sero-converted with a dose of 5×10^3^ IU (approximately 100,000-fold below a typical human adenovirus-vectored vaccine dose). These data suggest that the use of squalene oil-in-water emulsions with this vaccine- or indeed other adenovirus-vectored vaccines-may achieve either a modest increase in immunogenicity at high dose, or perhaps more likely a substantial dose-sparing and hence cost-reducing effect.

**Figure 4:**
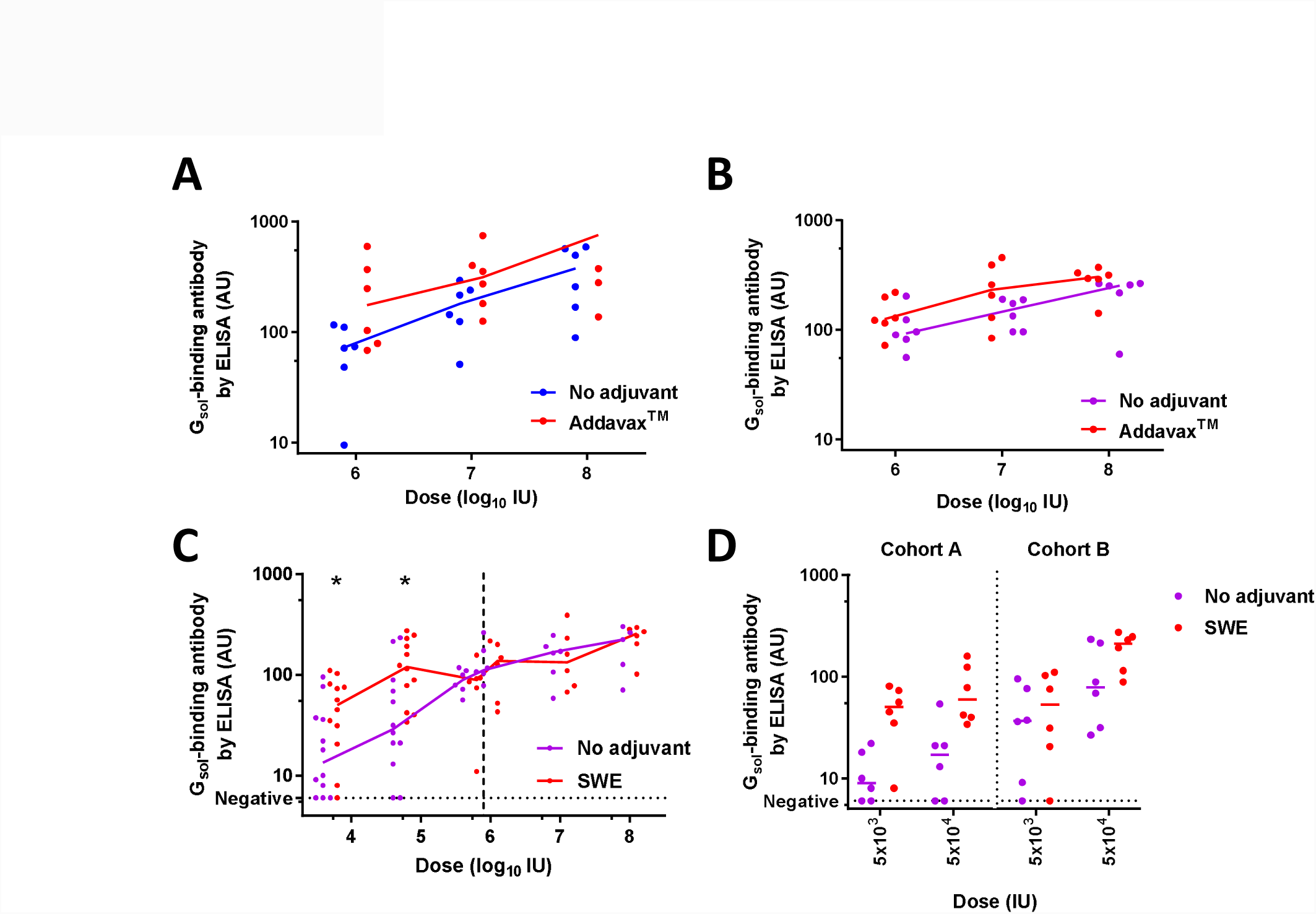
Adjuvantation of ChAdOx1 RabG and ChAdOX2 RabG. Panel A shows effect upon ELISA-measured antibody responses of addition of AddavaX™ to ChAdOx1 RabG. p=0.02 for effect of adjuvant by 2-way ANOVA performed upon log_10_-transformed data. Panel B shows effect upon ELISA-measured antibody responses of addition of Addavax™ to ChAdOX2 RabG. Data for mice not receiving the adjuvant is as shown for ChAdOX2 in Figure 3A. p=0.03 for effect of adjuvant by 2-way ANOVA performed upon log10-transformed data. Panel C shows effect upon ELISA-measured antibody responses of addition of SWE to ChAdOX2 RabG. Data shown combines results from three mouse cohorts: cohort A at doses 5×10^3^, 5×10^4^ and 5×10^5^ IU; cohort B at doses 5×10^3^ and 5×10^4^ IU; cohort C at doses 1×10^6^ to 1×10^8^ IU. Each experiment included 6 mice at each dose level, hence 12 individual results are shown for doses 5×10^3^ and 5×10^4^ IU. Stars indicate significant effect of adjuvant at a single dose level, pooling all data (p=0.05 at dose = 5×10^3^ IU, p=0.01 at dose = 5×10^4^ IU, both by two-tailed Mann-Whitney test). 2-way ANOVA was not performed across the full dose-range in view of the lack of adjuvant effect at high dose (i.e. a dose-adjuvant statistical interaction). Panel D shows the data from all mice receiving the 5×10^3^ and 5×10^4^ IU doses in cohorts A and B. This is the same data shown in panel C, but separating the two cohorts to show consistency of effect across experiments. P=0.0002 for effect of adjuvant by 3-way ANOVA performed upon log10-transformed data, with P=0.001 for effect of dose and P=0.0006 for effect of experiment; in keeping with the consistent trend of effect of adjuvant between doses and experiments, no statistical interaction between parameters was observed.

Throughout, points indicate results from individual mice; lines indicate group medians. All results shown are from serum samples collected 4 weeks after immunization. Units are arbitrary.

### ChAdOx2 RabG can be thermostabilized

We have previously described a simple method for thermostabilization of adenovirus-vectored vaccines by formulating the virus in a disaccharide-based solution and drying it onto a fibrous pad (40, 50). The materials required for this SMT technique are inexpensive (<USD 0.10 per dose) and the method is suitable for adaptation to GMP production. In-process losses of viral infectivity are close to zero. Other reported approaches to adenoviral thermostabilization include optimisation of liquid buffers (51, 52), lyophilization (53) and spray-drying (54, 55). To our knowledge, SMT is the only method reported to achieve adenoviral stability at temperatures in excess of 40 °C, such as may be encountered during ambient-temperature distribution in some low-income countries.

The SMT technique has not previously been applied to AdC68 or ChAdOx2 vectors. Here, we tested the ability of the technique to stabilize our ChAdOx2 RabG vector. Water content of the vitrified sugar-glass in the SMT product was 3.7% (median, n=3, range 3.4-3.8%). We observed excellent stability over 1 month at 45 °C, considerably out-performing virus formulated in the liquid buffer A438 (41): median log10 IU titer loss was 0.4 in the SMT product (Figure 5). No viable virus remained detectable in the A438. We are not aware of any other liquid buffer formulation which out-performs A438 for the stabilization of species E adenoviruses such as AdC68/ChAdOx2.

**Figure 5:**
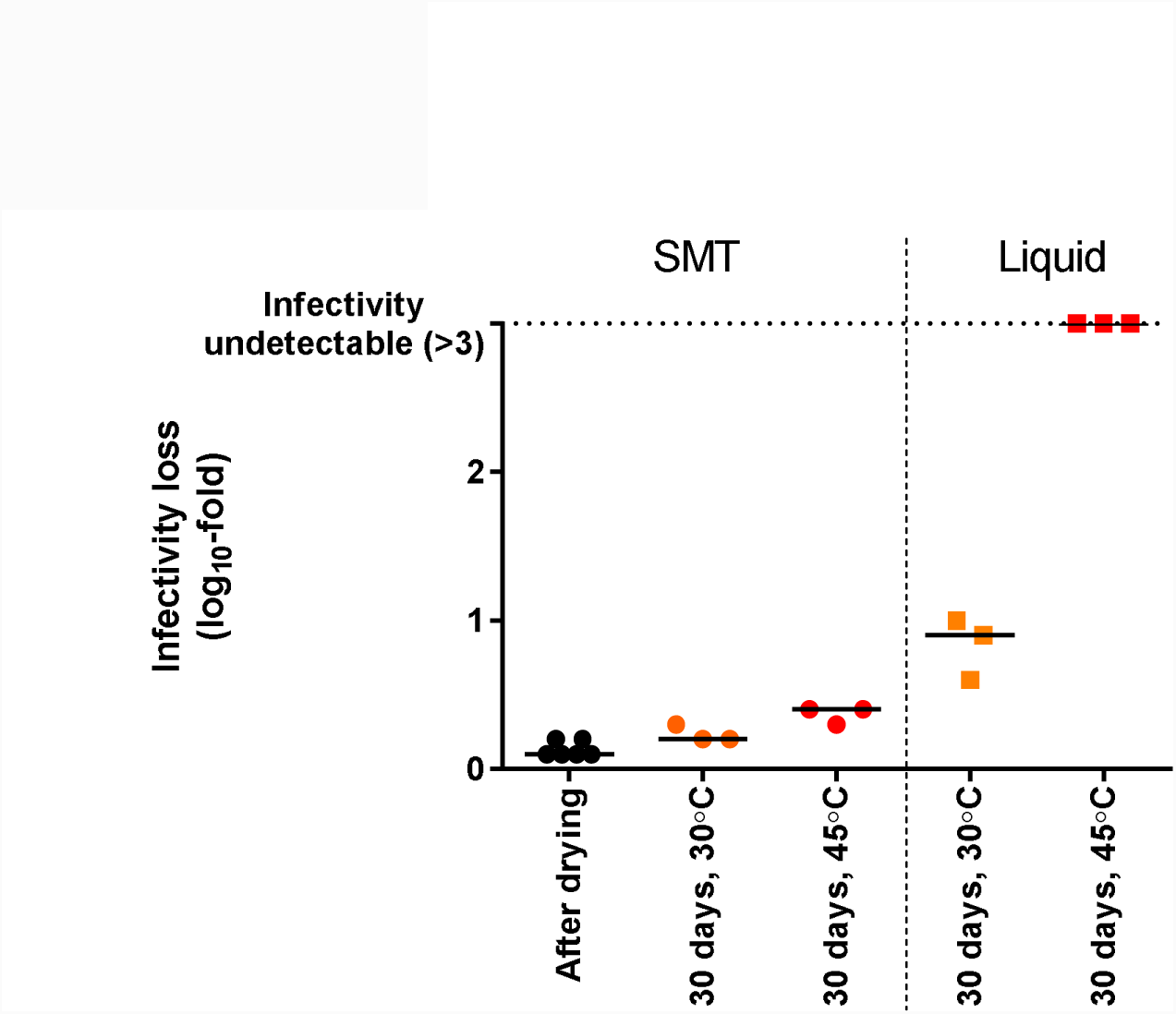
Sugar-matrix thermostabilization of ChAdOx2 RabG. ChAdOX2 RabG was dried using the SMT method or formulated in A438 liquid buffer before being stored at the indicated temperature for 30 days. IU titer was measured after drying (for SMT formulation) or after storage. Titer loss during storage was calculated by comparison to the titer of virus stored at −80 °C (in liquid formulation). Points indicate independent samples (i.e. separate vials); line indicates median for each condition.

## Conclusions

Global rabies mortality remains unacceptable for a disease which is technically straightforward to prevent-there is no immunological mystery regarding how to achieve protection against rabies. The obstacles to its control are primarily practical and economic. It is clear that increased effort and investment in existing methods of canine rabies control is both necessary and likely to prove productive. Nonetheless, the fact that rabies disease is concentrated in the least developed parts of the least developed countries poses significant challenges. In some such settings, there may be a role for pre-exposure vaccination of children against rabies, enabled by established infrastructure for the delivery of other childhood immunizations. We have therefore aimed to develop a technology which may render such an approach practical and cost-effective.

Here, we have built upon recent encouraging data with a closely related vaccine (23) to develop an iteratively improved candidate, ChAdOx2 RabG. We have shown that this has suitable genetic stability for GMP manufacture (Figure 1 and Results) and have preliminary indications of yield favourably comparable to other adenoviruses reported in the literature (46). Scalable adenovirus manufacturing platforms capable of producing virus titers exceeding 1×10^12^ VP per mL of culture have been developed (22, 56); this corresponds to a yield of 20 doses per mL at the typically-used human dose of 5×10^10^ VP, or 1000 doses per mL at the dose of 1×10^9^ VP at which AdC68.rabgp was protective in macaques (23). Such productivity is likely to be compatible with low-cost manufacturing.

In mice, a single dose of ChAdOx2 RabG reliably induced rabies virus neutralizing antibody (Figure 3). Responses to ChAdOx2 RabG compared favourably to those to AdC68.010 rabgp, the latter being closely related to the vector which achieved robust protection in a macaque-rabies challenge study (23). It is unclear why ChAdOx2 RabG appeared to outperform AdC68.010 rabgp at low vector doses; of the differences between the vectors, the one most likely to explain differing immunogenicity (as opposed to differing manufacturing characteristics) is the altered (intron-A containing) promoter used in ChAdOx2 RabG, which has previously been shown to enhance immunogenicity of other adenoviruses (27).

Our hope is that the ability of adenoviruses to achieve reliable, single-dose seroconversion in humans (57, 58) will enable ChAdOx2 RabG to achieve similar results in clinical trials, and hence to offer a single-dose alternative to existing PrEP regimes. We note though that the relatively slow kinetic of acquisition of antibody responses after ChAdOx2 RabG immunization (Figure 2B) may make the candidate less suitable for use in the PEP context, when urgent seroconversion is needed; indeed results were disappointing when AdC68.rabgp was used for PEP in non-human primates (23).

We were interested to observe that the relationship between ELISA-measured antibody and VNA titers differed for adenovirus-vectored and IRV vaccines (Figure 2D). Importantly, satisfactory VNA activity was achieved by the adenovirus-vectored vaccines (Figure 2C). Nonetheless, this altered relationship suggests that the adenovirus-vectored vaccines may be inducing a considerable amount of antibody which is detected by ELISA (i.e. binds soluble glycoprotein) but which does not neutralize the virus. Secreted soluble G (lacking the transmembrane domain) is known to adopt a conformation differing from that of the pre-fusion native G trimer, notably in that soluble G is predominantly monomeric (59). The antigen expressed by the adenovirus vectors is the full length (transmembrane-domain-containing) glycoprotein and would therefore be expected to form trimers on the surface of vector-infected cells. The epitopes displayed to B cells by such cells are still likely to differ from those displayed by rabies virions, not least in their spatial arrangement and, perhaps, steric accessibility of membrane-proximal regions. This observation suggests that, despite the good VNA results achieved with the current adenovirus-vectored vaccines, there may yet be scope for improvement in their efficacy, for example by engineering of the expressed antigen to focus the B cell response towards neutralizing epitopes.

We are encouraged by the observation that a dose-sparing effect may be achievable by formulating ChAdOx2 RabG with a squalene oil-in-water emulsion. Such adjuvants have an excellent clinical safety record and there are no longer intellectual property barriers to their use in most countries (49, 60)., Their raw materials cost cents per dose; manufacturing processes are published and sufficiently simple for transfer to academic or small commercial manufacturing organisations (34). The effect we have observed in mice is modest yet statistically significant and consistent across experiments with different vectors. Assessing whether such an effect is seen in humans would be relatively simple and low in cost, given access to existing GMP-grade adjuvant and adenovirus-vectored vaccine. Attainment of a 5 or 10-fold virus-dose-sparing effect using an inexpensive adjuvant could make a significant difference to the economic case for adoption of this or other adenovirus-based vaccines in low-resource settings.

As well as manufacturing cost, a further obstacle to vaccine delivery in such settings is the maintenance of a cold chain. This is particularly pertinent when considering the deployment of a live (albeit replication-deficient) viral-vectored vaccine. We have therefore sought to ‘build in’ thermostability from the early stages of development of the ChAdOx2 RabG candidate. Here, we demonstrated stabilization of ChAdOx2 resulting in ability to withstand 45 °C for 1 month with modest infectivity loss (0.4 log10-fold). This modest loss would be expected to have minimal effect upon immunogenicity (eg see the shallow dose-response curves in Figure 3A), but does suggest caution regarding longer-term storage at such temperatures. In its current form, SMT is probably adequate to allow adenovirus-vectored vaccines to meet the requirements for distribution via the WHO’s ‘controlled temperature chain’ (CTC) programme (i.e. maintenance of compliance with product specification after a single exposure to at least 40 °C for a minimum of 3 days just prior to administration) and/or use with chromogenic vaccine vial monitors (61, 62). There remains scope for further improvement of the SMT approach to enable true long-term ambient-temperature storage of such vaccines: we are currently pursuing a number of optimization strategies.

We have recently secured funding for GMP bio-manufacture and Phase I clinical trial of ChAdOx2 RabG in both liquid and SMT formulations. We hope ChAdOx2 RabG may eventually enable low-cost rabies PrEP in low-income settings. It may also have a role in PrEP for travellers to such settings (potentially facilitating clinical development by providing a ‘dual market’ in high-income as well as low/middle-income countries). The SMT technology is applicable to multiple adenovirus-vectored vaccines and other vaccine platforms (40, 50); we thus hope that, as well as developing a tool with potential for rabies control, this trial will advance a technology with broad utility for the delivery of vaccination to the millions of individuals who live in resource-poor settings.

## Acknowledgements

We are grateful for the assistance of the Jenner Institute Viral Vector Core Facility in the production of adenoviruses, for assistance with ELISA from Kunitaka Yoshida, and for critical review of the manuscript by Adam Ritchie.

Author contributions
Conceptualization, funding acquisition, project administration, supervision: ADD, RA, HCE, AVSH, SJD, ARF; Investigation: CW, PD, XZ, ZX, HG, AB, ADD; Resources: LB, RV, NC; Writing-original: ADD; Writing-review and editing: All

